# Bottom trawl catch comparison in the Mediterranean Sea: Flexible Turtle Excluder Device (TED) *vs* traditional gear

**DOI:** 10.1101/610089

**Authors:** Claudio Vasapollo, Massimo Virgili, Andrea Petetta, Giada Bargione, Antonello Sala, Alessandro Lucchetti

**Affiliations:** National Research Council (CNR), Institute of Biological Resources and Marine Biotechnologies (IRBIM) of Ancona, Largo Fiera della Pesca, 1 – 60125 Ancona, Italy; Department of Biological, Geological and environmental sciences, University of Bologna, Piazza di Porta San Donato 1 – 40126 Bologna, Italy

## Abstract

The Mediterranean Sea is a hotspot of biodiversity, but the high fishing pressure results in high bycatch rates of protected (sea turtles and cetaceans) and top predator species (sharks). The reduction of bycatch is challenging for fishery scientists, as conservation of these species has become a priority. Among the animals threatened by fishing activities, the loggerhead (*Caretta caretta*) represents a charismatic species considered as “vulnerable” at the global scale by IUCN.

In the Mediterranean Sea, trawl nets show the highest probabilities of bycatch of protected species, with high rates of mortality. A new flexible Turtle Excluder Device (TED) has been tested for the first time on a commercial scale in the Mediterranean Sea to assess its effectiveness in reducing bycatch. The results did not show any significant (α = 0.05) loss in terms of commercial weight, but a significant reduction of debris in the codend of the nets mounting the TED respect to traditional nets. The catch comparison of the main commercial species showed similar rates without any significant loss of sizes, with the only exception of anglerfishes (*Lophius* spp.) that showed a loss of the largest individuals by TED. In terms of bycatch, the traditional nets captured mostly rays and sharks, while no turtles were captured, at all. In this regard, the authors were informed by other vessels operating in the same areas at the time of the trials about some accidental catches of loggerhead turtles. Our results demonstrated that the flexible TED represents a practical and effective solution to reduce the bycatch of endangered species in coastal Mediterranean demersal multispecies fisheries, as demonstrated experimentally also in other areas of the world. The measures involving technical modifications of fishing gears require significant investments but are technically feasible and could guarantee the success of the conservation.

## Introduction

The Mediterranean fishing fleet is highly diversified and targeting several species. The Mediterranean basin is also considered as a hot spot of biodiversity [1]. Nevertheless, the high fishing effort has resulted in overexploitation of fish resources [2] and in the deterioration of marine ecosystem services [3,4]. The bycatch of protected species, such as cetaceans [5] and sea turtles [6] and top predator species, such as sharks without any economic importance [7] is a consequence of the intense fishing pressure. The solution to reduce the bycatch has become a challenge for fishery scientists as the conservation of species has become a priority for large international organizations. The Habitats Directive [8], for examples, imposed, among others, a conservation policy aimed at reducing the bycatch of species present in the list animals of Annex IV. FAO’s International Guidelines on Bycatch Management and Reduction of Discards [9] identified management measures necessary to ensure the conservation of target and non-target species. Conservation of the megafauna is an intricateed link of multiple ocean resources that act in a dynamic and complex ecological ocean system that varies in a wide range of spatial and temporal scales. To make matters worse, conservation is often further complicated by competing factors such as social, economic and ecological and related management objectives [10].

Among the animals threatened by fishing activities, the loggerhead sea turtle (*Caretta caretta*) is a charismatic species considered as “vulnerable” at the global scale [11] and as “least concern” for the Mediterranean sea [12].

Nevertheless, the adoption of conservation actions for the species is still a crucial point in the Mediterranean. Due to their habits, such as breeding and feeding migrations, loggerhead turtles interact with several types of fishing gears (towed gears, set nets and longlines) [13,14]. In [15] the author estimated that more than 150000 captures per year occur in the Mediterranean due to fishery, with more than 50000 deaths per year. [13] estimated that around the Italian coasts more than 52000 turtles are caught per year with a mortality of 10000 individuals.

Among the fishing gears, trawl nets showed the highest probabilities of bycatch, thus representing the most dangerous in terms of mortality for loggerhead [16]. A situation of great concern is particularly strong in the Adriatic Sea, where due to its shallow waters, representing a favourable fishing ground, is exploited by more than 1000 bottom trawlers, owning mainly to the Italian and Croatian commercial fleets [14]. This area also represents a suitable foraging site for sea turtles [17], where they can find rich benthic communities to feed on. Due to the massive presence of fishing vessels and loggerhead individuals, the northwestern Adriatic is considered as a bycatch hotspot where the probability of encounter between a trawler and a turtle is considerably high above all in late summer-autumn [16]. The annual trawlers bycatch of sea turtles in the northern Adriatic was estimated as more than 6500 individuals [13,18].

A specific technical measure was proposed in the late 1980s to reduce the sea turtle mortality, i.e., the Turtle Excluder Device (TED) [19,20]. TED is a rounded grid, which stops large objects or animals and expels them by an exit placed before the codend. TEDs showed their effectiveness mainly in prawn trawl fisheries, to the point that in several countries the use of TED is mandatory [14]. Nevertheless, [18] were skeptical about the TED efficency in the Mediterranean waters because they would exclude the larger commercial individuals too. On the other hand, recent experiments held in the Mediterranean sea [21,22] did not show any loss in term of the commercial catch. During those scientific trawl surveys they obtained as a result that the main commercial species did not show any decrease in terms of weight and number of individuals respect to a net without the grid. Moreover, also a decrease of debris was observed, which translates to improved catch quality and a reduction of additional sorting operations on-board, increasing fishing time and earn.

With these premises, the aim of the present paper is to assess the effectiveness of a TED on commercial scale in different areas of the northern Adriatic targeting different commercial species. The objectives were: 1) to study the gear performance in the presence of TED, 2) to compare the catch rates of commercial species, as well as of the discarded species and debris (both anthropogenic and natural) and 3) to analyse the eventual size selection induced by the presence of the TED using a length-based analysis of the main commercial species.

## Materials and methods

### Sea trials

Seven bottom trawlers were randomly selected from different harbours from the northern to the central Italian coasts to conduct the sea trials (Fig 1). Two of the boats were twin trawlers; in this case, one net was armed with the TED and the other not (called control, CTRL). The rest of the boats were single traditional net trawlers. For these latter, two cruises per boat were needed: one mounting the TED and the other without. The boats were coded based on the first three letters of the boat names as follows: AUD = Audace (twin-trawler), RIM = Rimas, JOA = Joachì, AST = Astuzia, GLA = Gladiatore (twin-trawler), PAL = Palestini and TAR = Tarantini. Trials were made in 2015, 2016 and 2017 between June and December (Table 1). The boat crews made fishing operations during normal commercial fishing activities, while scientific observers on-board collected measures and weighs.

**Table 1.** Main characteristics of the boats used for the trials. AUD: Audace, RIM: Rimas, JOA: Joacchì, AST: Astuzia, GLA: Gladiatore, PAL: Palestini, TAR: Tarantini. LOA: Length overall, GT: Gross Tonnage.

| Boat | Power<br>[kwh] | LOA<br>[m] | GT<br>[tonns] | Month of<br>sampling | Year of<br>sampling |
| --- | --- | --- | --- | --- | --- |
| AUD | 872 | 26.8 | 130 | July | 2015 |
| RIM | 142 | 13.6 | 15 | November | 2015 |
| JOA | 574 | 26.2 | 96 | March | 2016 |
| AST | 147 | 15.5 | 16 | June | 2016 |
| GLA | 870 | 26.5 | 96 | December | 2016 |
| PAL | 206 | 22.3 | 50 | April | 2017 |
| TAR | 167.9 | 17.3 | 24.4 | July | 2017 |

**Fig. 1.**
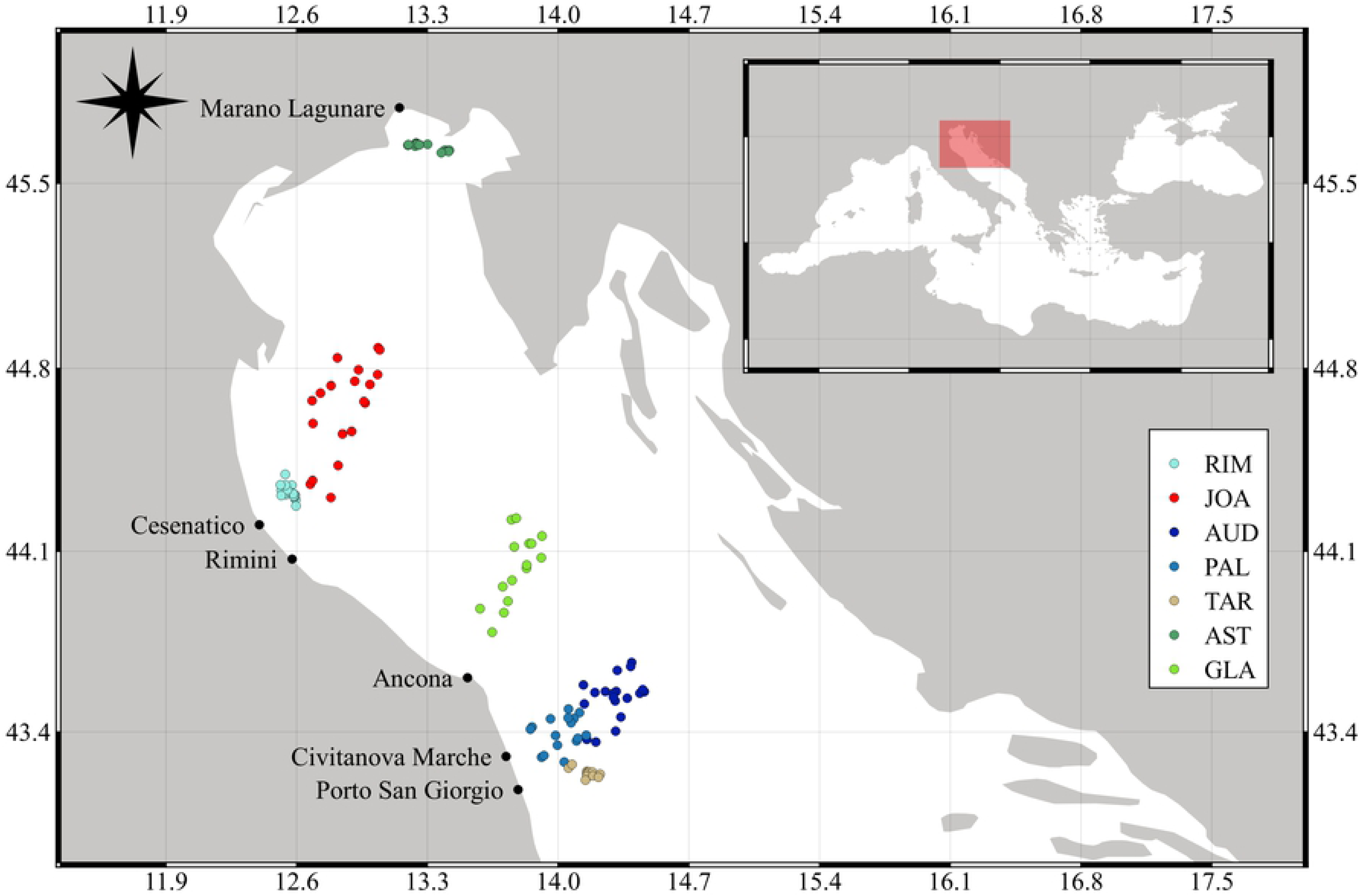
The study area and hauls made during sea trials. The base harbour for each boat are reported. AUD: Audace, RIM: Rimas, JOA: Joacchì, AST: Astuzia, GLA: Gladiatore, PAL: Palestini, TAR: Tarantini.

### TED specifications

The flexible TED used for the trials, was made of an alloy of high strength plastic material. It was designed according to the technical specifications suggested by Mitchell et al. (1995) (Fig 2). The TED was mounted on a tubular netting section (6 m in length) with a tilt angle of approximately 46°, and placed in the extension piece, just in front of the codend of the commercial trawl nets. An escape opening was cut on the upper portion of the net just before the TED and covered by a netting panel with three sides sewn to the net to prevent loss of commercial species. This panel operated like a valve, as it opened only when it was hit by large and heavy objects, and thus allowing sea turtles and other bycatch species to out the net. According to [23], an accelerator funnel was installed before the TED for driving fishes down and away from the exit panel and through the TED bars, toward the codend.

**Fig. 2.**
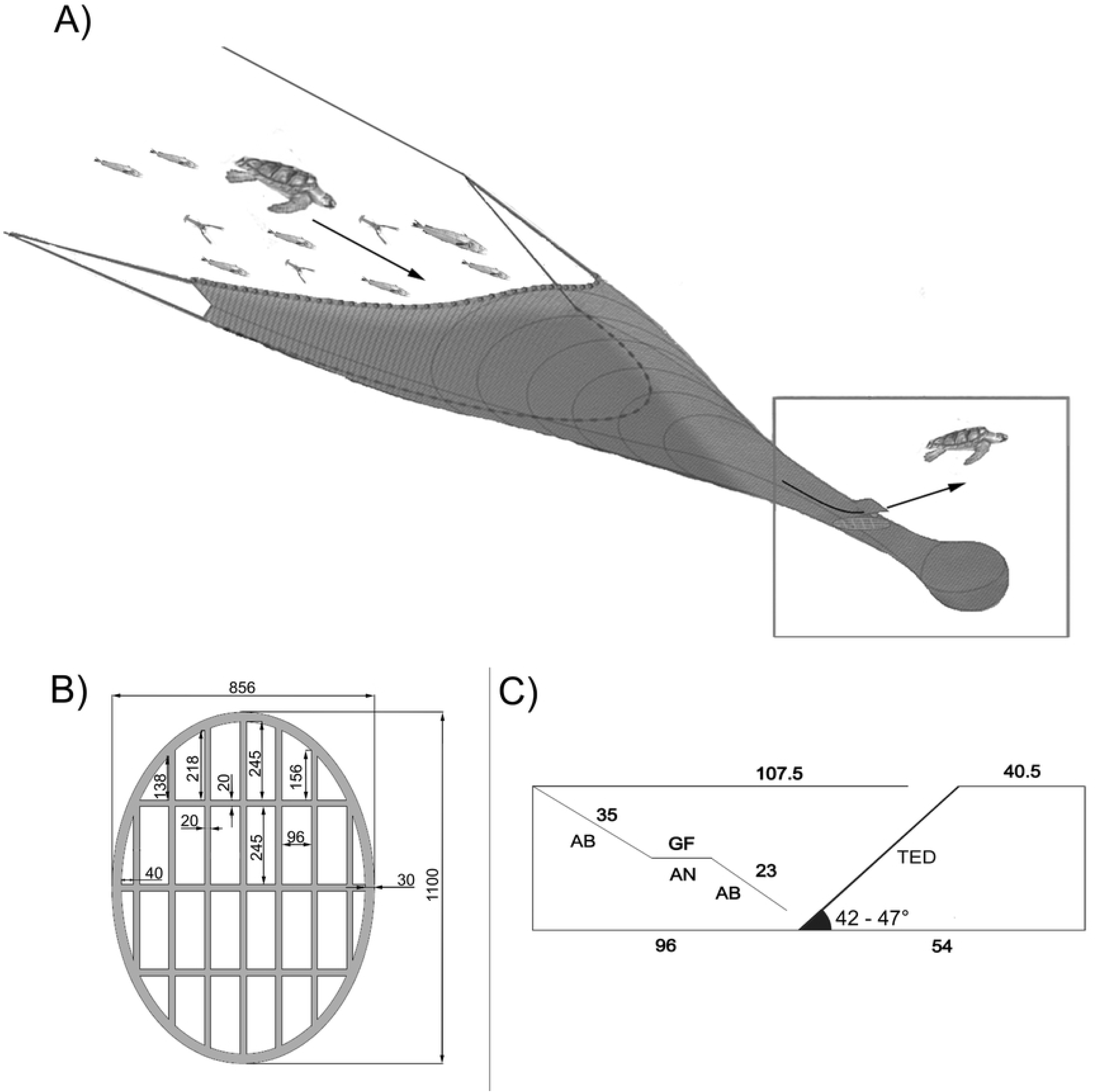
Flexible Turtle Excluder Device scheme. a) representation of the position of the TED in relation to the codend. b) design of the flexible TED used during the sea trials. Size in mm. c) Technical drawing of the TED rigging (lateral view). AB and AN, types of net cuttings (figures indicate number of meshes). The average grid angle recorded during sampling is also reported.

The TED angle is a key factor influencing TED efficiency and preventing loss of commercial species during the tow [23,24]. An angle less than 40° may involve catch loss due to water diversion through the exit hole. Angles greater than 55° can prevent turtle escape and deflection of trash, clogging the grid. Therefore, TED performance was measured using the Star Oddi Data Storage Tags (DST) sensors (Iceland) to assess the grid’s angle. Sensors were directly mounted on the grid and sampled information on TED angle every 60 s.

TED performance, fish reaction to the TED and fish behaviour inside the net were also monitored using an underwater camera (GoPro Hero4, US). Due to high water turbidity, the camera was mounted 1 m from the TED. The fisheye consistently provided a full view of the TED monitoring grid position during hauling.

### Catch analysis

Catches for each hauls were subdivided into three categories: commercial (including commercially important species), discard (including those species, both invertebrates and fishes not commercially important or under legal size, if any) and debris (including material, both anthropogenic and natural, like stones and woods, that is considered as litter).

Catches were standardized based on the formulas:

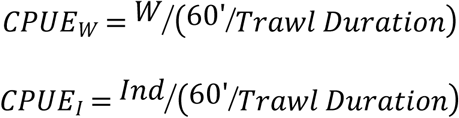

where *CPUE*_*W*_ is the catch per unit effort expressed in terms of weight (kg) per hour of trawling and *CPUE*_*I*_ is the catch in terms of individuals caught per hour, *W* is the weight of the catch of each single haul and the *Trawl Duration* is the time the net fished in each single haul expressed in minutes. Wilcoxon’s Rank Test was used to assess if differences emerged between the catches of the nets with and without TED for each boat, as well as tow duration and depth of sampling [25].

To compare the catches as commercial CPUE_W_, as well as discards and debris, a Generalized Linear Mixed Model (GLMM) was used. The independent variables net (TED *vs* CTRL), depth (coded as “Low”, between 11 and 30 m; “Medium”, between 30 and 50 m; “High”, between 50 and 88 m) and year were at first tested for co-linearity both visually (with a scatter plot of each variable *vs* each other) and by Pearson’s correlations. The boat term was considered as random factor. The model selection was made based on both the Akaike’s Information Criterion (AIC) and the Log Likelihood Ratio Test, following the protocol in [26]. Any residuals trends and heteroscedasticity were assessed to check if statistical assumptions were respected. If variance heterogeneity was observed associated to a variable, the variance structure of the model was modified to allow a different variance for each level of the variable [26,27]. When one or more factors of the models resulted as significant (p < 0.05) a pairwise test based on Tukey’s test was adopted to investigate which are the levels that showed significantly different mean values.

For commercial species, the total length (TL) of each specimen was measured on-board the vessels to the nearest 0.5 cm below. To assess the influence of the TED on the size of the fish caught, the length frequency distributions (LFD) for the commercial species representing more than 5% of the total catch in weight for each boat were analysed. The catch comparison to apprise the catch efficiency (at length) of TED relative to CTRL was made using GLMM. The probability of a fish being retained by TED follows from:

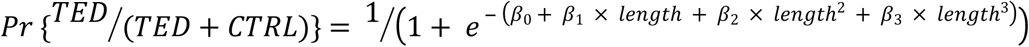

A binomial error distribution was used to calculate the probability of the number of fish caught in the TED gear given they enter both gears by 1-cm size class. A probability value of 0.5 corresponds to equal catches in both gears. According to [28] a 3^rd^ order polynomial would be adequate for most cases, although in some instances a 1^st^ or 2^nd^ order would be enough. The best binomial model was chosen based on AIC. A random term was added to the models. In papers aimed at testing a gear relative to another, paired hauls were analysed considering hauls as random effects [28–32]. In our case, as for some boats the hauls were not paired between TED and CTRL, the hauls were pooled together for each boat, and the term boat was used as a random intercept, instead. Moreover, as the individual lengths of some species were not always the same among the boats, length was used as a random slope. The species selected correspond to the target species of the period the boats were fishing, thus not all the vessels caught the same species in the same proportion. Consequently, the models for each species run with a different number of vessels. The models are reported with a 95% confidence interval calculated with a bootstrap method using 999 simulations.

Any sea turtle eventually caught (as well as other bycatch species) were measured (curved carapace length, CCL, in cm) and weighed, and then rescued.

All the analyses were performed using the free software R [33] and the R packages *nlme* [34] and *lme4* [35].

## Results

Overall, 153 hauls were made for comparing the efficiency of TED *vs* CTRL net (Table 2). Wilcoxon’s test did not show any differences in the tow duration between TED and CTRL trawls for any of the boats, apart for TAR that showed a marginal significant difference of about 6 minutes (Table 2). Also for average fishing depth, no differences were observed with the only exception of JOA, but this could probably be the effect of the unbalanced number of hauls between TED and CTRL.

**Table 2.** Operating conditions and catch per unit effort based on weight (kg h^−1^) for each boat during the sea trials. AVG: Average (per haul), SE: standard error, ns: not significant. P value refers to the Wilcoxon’s test statistic. AUD: Audace, RIM: Rimas, JOA: Joacchì, AST: Astuzia, GLA: Gladiatore, PAL: Palestini, TAR: Tarantini.

|  |  |  | Tilt<br>[degree] | Angle | Tow<br>[min] | Duration |  |  | Depth<br>[m] |  |  |  | Commercial<br>[kg h<sup>-1</sup>] |  | Catch |  | Discard<br>[kg h<sup>-1</sup>] |  | Catch |  | Debris<br>[kg h<sup>-1</sup>] |  | Catch |  |
| --- | --- | --- | --- | --- | --- | --- | --- | --- | --- | --- | --- | --- | --- | --- | --- | --- | --- | --- | --- | --- | --- | --- | --- | --- |
| Boat | Net<br>Type | No.<br>Hauls | AVG | SE | AVG | SE | Range | p value | AVG | SE | Range | p value | AVG | SE | Range | p value | AVG | SE | Range | p value | AVG | SE | Range | p value |
| AUD | TED | 18 | 46.0 | 0.7 | 139.2 | 1.0 | 97-169 | ns | 78.9 | 0.3 | 67.3-88.0 | ns | 10.2 | 0.7 | 5.4-15.3 | ns | 9.6 | 0.9 | 4.4-18.4 | << 0.001 | 5.3 | 0.6 | 1.7-10.8 | << 0.001 |
|  | CTRL | 18 |  |  | 139.2 | 1.0 | 97-169 |  | 78.9 | 0.3 | 67.3-88.0 |  | 13.3 | 1.2 | 7.4-28.7 |  | 12.8 | 1.4 | 3.7-24.1 |  | 10.6 | 1.1 | 4.2-21.1 |  |
| RIM | TED | 10 | 44.9 | 0.9 | 139.4 | 1.2 | 122-166 | ns | 18.7 | 0.3 | 12.1-23.7 | ns | 40.0 | 2.3 | 28.4-50.7 | ns | 17.7 | 1.5 | 12.3-26.7 | ns | 3.8 | 0.6 | 1.7-7.3 | ns |
|  | CTRL | 10 |  |  | 143.2 | 1.3 | 113-167 |  | 18.2 | 0.2 | 13.0-21.6 |  | 40.3 | 3.2 | 24.6-56.4 |  | 15.5 | 2.0 | 7.8-29.8 |  | 5.7 | 1.1 | 1.3-10.1 |  |
| JOA | TED | 10 | 41.5 | 0.5 | 116.0 | 0.7 | 100-130 | ns | 34.1 | 0.4 | 20.0-38.3 | 0.033 | 13.7 | 1.1 | 8.8-19.9 | ns | 12.2 | 2.2 | 36.8-41.2 | ns | 4.5 | 1.1 | 0.7-9.9 | ns |
|  | CTRL | 9 |  |  | 109.1 | 0.7 | 95-130 |  | 30.6 | 0.2 | 25.0-35.0 |  | 14.6 | 0.7 | 10.7-17.4 |  | 11.7 | 20.6 | 4.4-30.4 |  | 4.0 | 1.3 | 0.5-12.1 |  |
| AST | TED | 10 | 42.4 | 0.6 | 66.1 | 1.3 | 50-100 | ns | 13.5 | 0.1 | 12.9-15.4 | ns | 12.2 | 1.9 | 4.7-24.9 | ns | 126.8 | 27.6 | 18.6-271.5 | ns | 36.6 | 9.9 | 5.4-109.7 | 0.048 |
|  | CTRL | 8 |  |  | 84.2 | 2.5 | 45-115 |  | 12.6 | 0.1 | 11.9-15.0 |  | 11.7 | 3.5 | 4.4-30.4 |  | 107.0 | 44.2 | 36.9-348.7 |  | 21.5 | 10.8 | 4.4-84.5 |  |
| GLA | TED | 15 | 47.1 | 0.7 | 117.3 | 0.5 | 95-132 | ns | 39.1 | 0.2 | 26.0-43.2 | ns | 21.6 | 1.4 | 12.5-32.5 | ns | 22.7 | 3.7 | 7.7-47.8 | 0.046 | 1.5 | 0.3 | 0.0-3.3 | 0.002 |
|  | CTRL | 15 |  |  | 117.3 | 0.5 | 95-133 |  | 39.1 | 0.2 | 26.0-43.2 |  | 25.0 | 1.3 | 16.2-35.0 |  | 31.3 | 4.6 | 7.8-59.1 |  | 5.2 | 1.0 | 1.6-17.3 |  |
| PAL | TED | 8 | 43.5 | 0.9 | 145.8 | 1.8 | 130-200 | ns | 49.3 | 0.9 | 21.4-61.0 | ns | 10.5 | 1.0 | 6.5-14.0 | ns | 5.5 | 0.8 | 3.1-8.9 | 0.018 | 8.5 | 0.9 | 3.9-12.7 | ns |
|  | CTRL | 8 |  |  | 133.9 | 0.6 | 130-150 |  | 46.3 | 1.3 | 23.0-66.0 |  | 7.4 | 0.8 | 4.5-11.1 |  | 8.6 | 1.6 | 10.7-15.6 |  | 10.0 | 1.8 | 1.8-16.6 |  |
| TAR | TED | 7 | 44.6 | 1.5 | 146.6 | 0.4 | 135-155 | 0.048 | 62.1 | 1.1 | 30.0-73.5 | ns | 12.4 | 0.8 | 10.1-15.9 | ns | 23.2 | 0.8 | 19.2-26.3 | ns | 9.7 | 1.4 | 4.6-13.9 | ns |
|  | CTRL | 7 |  |  | 140.2 | 0.3 | 135-146 |  | 61.9 | 1.1 | 34.6-74.3 |  | 13.1 | 0.6 | 10.7-15.6 |  | 17.3 | 2.2 | 12.3-126.5 |  | 15.9 | 2.6 | 9.5-29.7 |  |

### Gear performance

Images of underwater camera and sensor’s data showed, for all vessels, that the TED did not affect the functioning of the net. The tilt angle of the TED (Table 2), obtained from > 7200 pings (> 120 hours) on the whole boats, ranged, on average, between 41.5° ± 0.5° (mean ± SE, hereafter) and 47.1° ± 0.7°.

### Catch rates

The results from the catch averages are summarised in Table 2. Wilcoxson’s test for commercial standardized catch did not show any differences between TED and CTRL for any of the boats. Discards, on the other hand, showed significant differences for AUD, GLA (although negligible) and PAL, with TED always showing lower values than CTRL. For Debris also, three boats showed significant differences: AUD, AST (although negligible) and GLA. Again, the differences are in favour of TED showing less weight, apart for AST where TED appeared to have caught more than CTRL, but this difference is statistically borderline at α = 0.05. Lists of the commercial species, discards and debris categories are available as S1-S3 Tables.

The model selections to test the effects of the explanatory variables on CPUE_W_ are shown in Table 3.

**Table 3.**
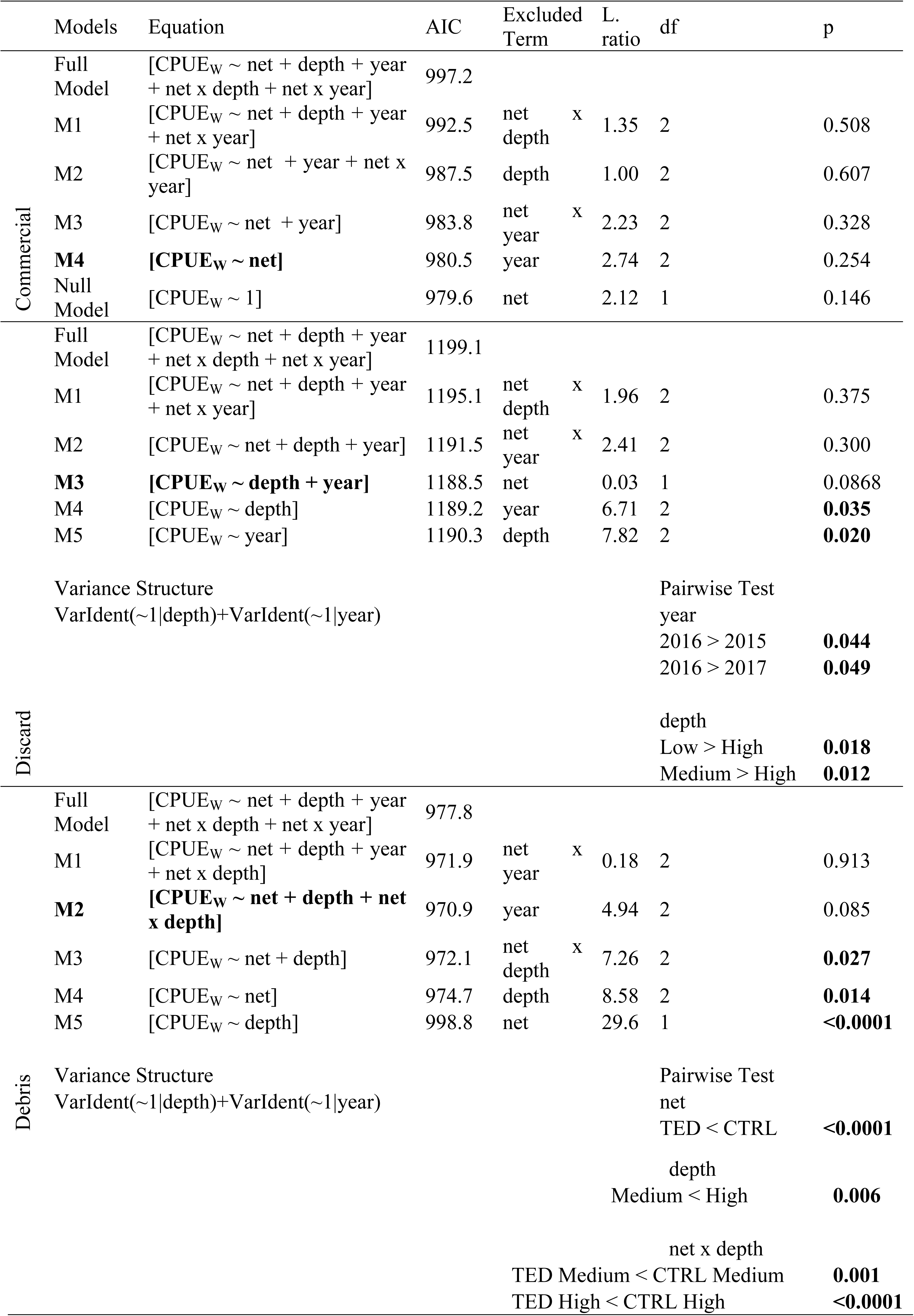
Model selection for the catch rates. In bold between square brackets the best model explained by the independent variables. In bold outside the square brackets the p value of the significant terms in the models.

No differences in total commercial catches were observed when adopting TED or not (17.1 ± 1.2 and 18.7 ± 1.4 CPUE_W_, for TED and CTRL, respectively; Fig 3). The best model for discard (Fig 3, Table 3) comprised only the depth and the year of sampling. The pairwise based on factor year showed that 2016 was the year when more discard was caught, but the significance is marginal. The pairwise for factor depth highlighted that more discard was present in low and medium depths respect to high depths (51.4 ± 10.3, 30.2 ± 5.4 and 12.2 ± 0.9 CPUE_W_, respectively). For debris (Fig 3) the best model comprised net, depth and their interactions (Table 3).TED CPUE_W_ was lower than CTRL (8.9 ± 1.8 and 9.5 ± 1.2 CPUE_W_, respectively) as stressed by the pairwise test. Differences exists also between medium and high depths (4.2 ± 0.5 and 9.5 ± 0.7 CPUE_W_, respectively). The pairwise for the interaction term showed differences between TED and CTRL at medium depth (3.2 ± 0.6 and 5.5 ± 0.8 CPUE_W_, respectively) and at high depth (6.9 ± 0.7 and 12.2 ± 1.1 CPUE_W_, respectively).

**Fig. 3.**
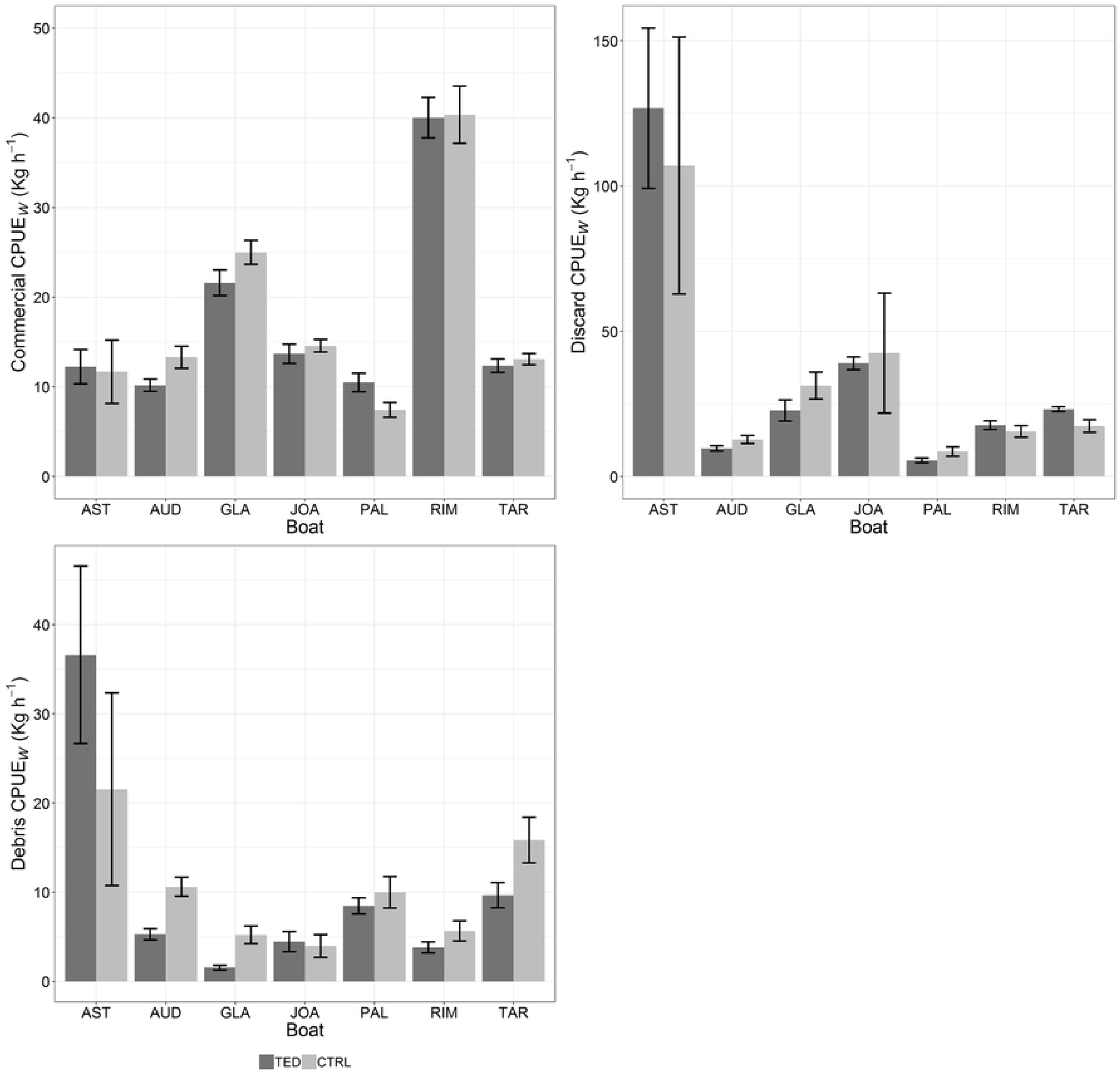
Average commercial, discard and debris CPUEs in weight for TED and CTRL nets per boat. Bars are standard errors. AUD: Audace, RIM: Rimas, JOA: Joacchì, AST: Astuzia, GLA: Gladiatore, PAL: Palestini, TAR: Tarantini.

Eight commercial species were selected that respected the 5% threshold of the total catch in weight in at least two boats, for catch comparison analysis: *Lophius* spp., *Merluccius merluccius* (minimum landing size, MLS, 20 cm), *Mullus barbatus* (MLS = 11 cm), *Illex coindetii, Sepia officinalis, Melicerthus kerathurus, Parapenaeus longirostris* (MLS = 20 mm) and *Squilla mantis*. LFDs are represented in Fig 4. It is evident how for the same species, the LFDs are different among boats, depending mostly on the area, on the period of fishing and, as well as, on the depth (as a proxy for the distance from the coast. When the individuals were pooled together for each single species (Fig 5), the differences between TED and CTRL, if any, were more evident. The parameter estimates for the fit of the proportion of individuals caught by TED respect to CTRL are detailed in Table 4. In Fig 6, the general trends of the proportion of individuals caught by TED and CTRL are shown together with trends for each single boat (images extrapolated by the videos showing small sized species are showed in S1 FIg). For the three fish species, TED appeared to be more efficient in catching small individuals, while increasing the fish length the proportion decreases, although for *M. merluccius* and for *M. barbatus* the ratio is almost near the value 0.5 indicating that both nets caught similar numbers of fishes. On the contrary, for *Lophius* spp. the ratio decreases in favour of CTRL when length increases, but it is noteworthy to consider that the bulk of the catch is comprised between 20 and 30 cm, with the longest fishes (> 30 cm) representing a small percentage of the total catch. Some concerns emerged with this species. *Lophius* spp. are characterized by a big head, which sometimes prevents the passage of the fish through the grid bars. Sometimes medium to large individual reached the TED transversely (S2 Fig), remaining mashed on the grid bars. In some instances, the individuals were pushed to enter the grid by the hydrodynamic force, while in other cases, the animals rolled up until to reach the opening on the upper side of the TED. Considering the two molluscs, *S. officinalis* showed a constant slope while increasing sizes and the ratio is always above the 0.5 value, but extremely near to it. For *I. coindetii*, after a constancy in the ratio near the 0.5 value, the trend decreases for larger animals. For *P. longirostris* the ratio is slightly lower than 0.5. Both for *M. kerathurus* and *S. mantis*, TED caught less individuals of small sizes respect to CTRL and, on the contrary, proportionally more of larger animals.

**Table 4.**
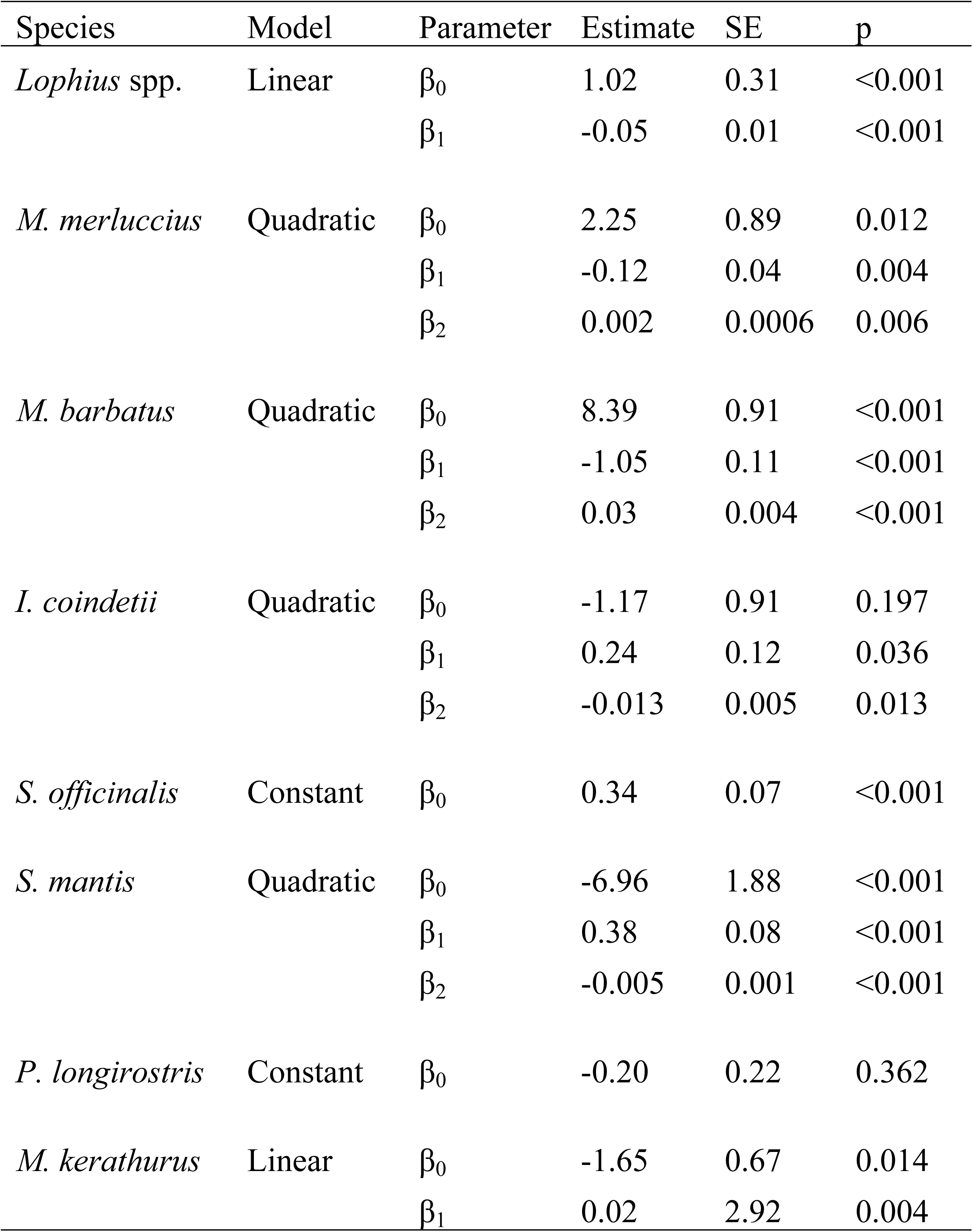
GLMM parameter estimates of the catch selectivity logistic models from the trials. SE: Standard Error.

| Species | Model | Parameter | Estimate | SE | p |
| --- | --- | --- | --- | --- | --- |
| *Lophius* spp. | Linear | $\beta_0$ | 1.02 | 0.31 | <0.001 |
| | | $\beta_1$ | -0.05 | 0.01 | <0.001 |
| *M. merluccius* | Quadratic | $\beta_0$ | 2.25 | 0.89 | 0.012 |
| | | $\beta_1$ | -0.12 | 0.04 | 0.004 |
| | | $\beta_2$ | 0.002 | 0.0006 | 0.006 |
| *M. barbatus* | Quadratic | $\beta_0$ | 8.39 | 0.91 | <0.001 |
| | | $\beta_1$ | -1.05 | 0.11 | <0.001 |
| | | $\beta_2$ | 0.03 | 0.004 | <0.001 |
| *I. coindetii* | Quadratic | $\beta_0$ | -1.17 | 0.91 | 0.197 |
| | | $\beta_1$ | 0.24 | 0.12 | 0.036 |
| | | $\beta_2$ | -0.013 | 0.005 | 0.013 |
| *S. officinalis* | Constant | $\beta_0$ | 0.34 | 0.07 | <0.001 |
| *S. mantis* | Quadratic | $\beta_0$ | -6.96 | 1.88 | <0.001 |
| | | $\beta_1$ | 0.38 | 0.08 | <0.001 |
| | | $\beta_2$ | -0.005 | 0.001 | <0.001 |
| *P. longirostris* | Constant | $\beta_0$ | -0.20 | 0.22 | 0.362 |
| *M. kerathurus* | Linear | $\beta_0$ | -1.65 | 0.67 | 0.014 |
| | | $\beta_1$ | 0.02 | 2.92 | 0.004 |

**Fig. 4.**
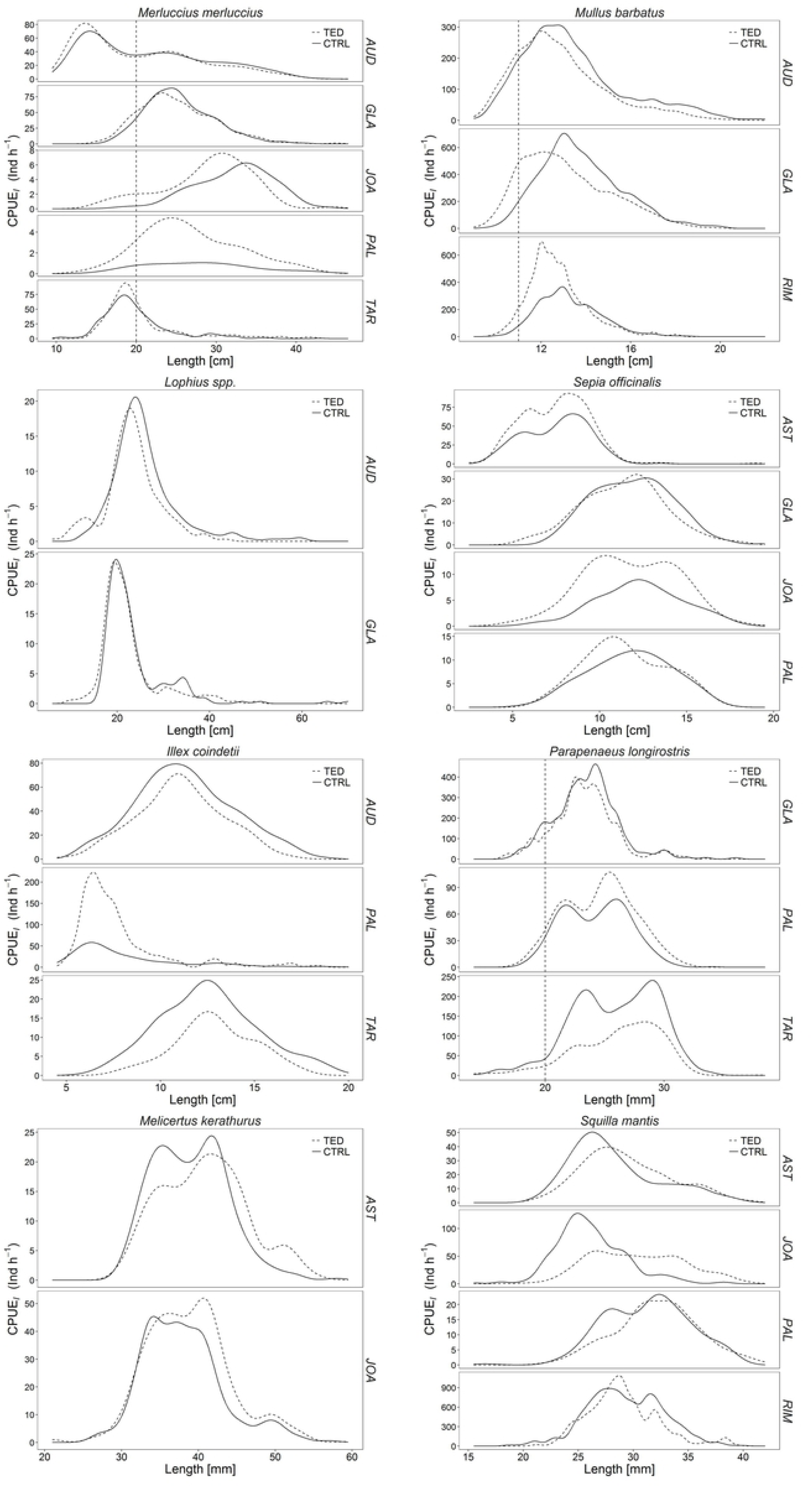
Catch length-frequency distributions for each commercially important species per boat. AUD: Audace, RIM: Rimas, JOA: Joacchì, AST: Astuzia, GLA: Gladiatore, PAL: Palestini, TAR: Tarantini.

**Fig. 5.**
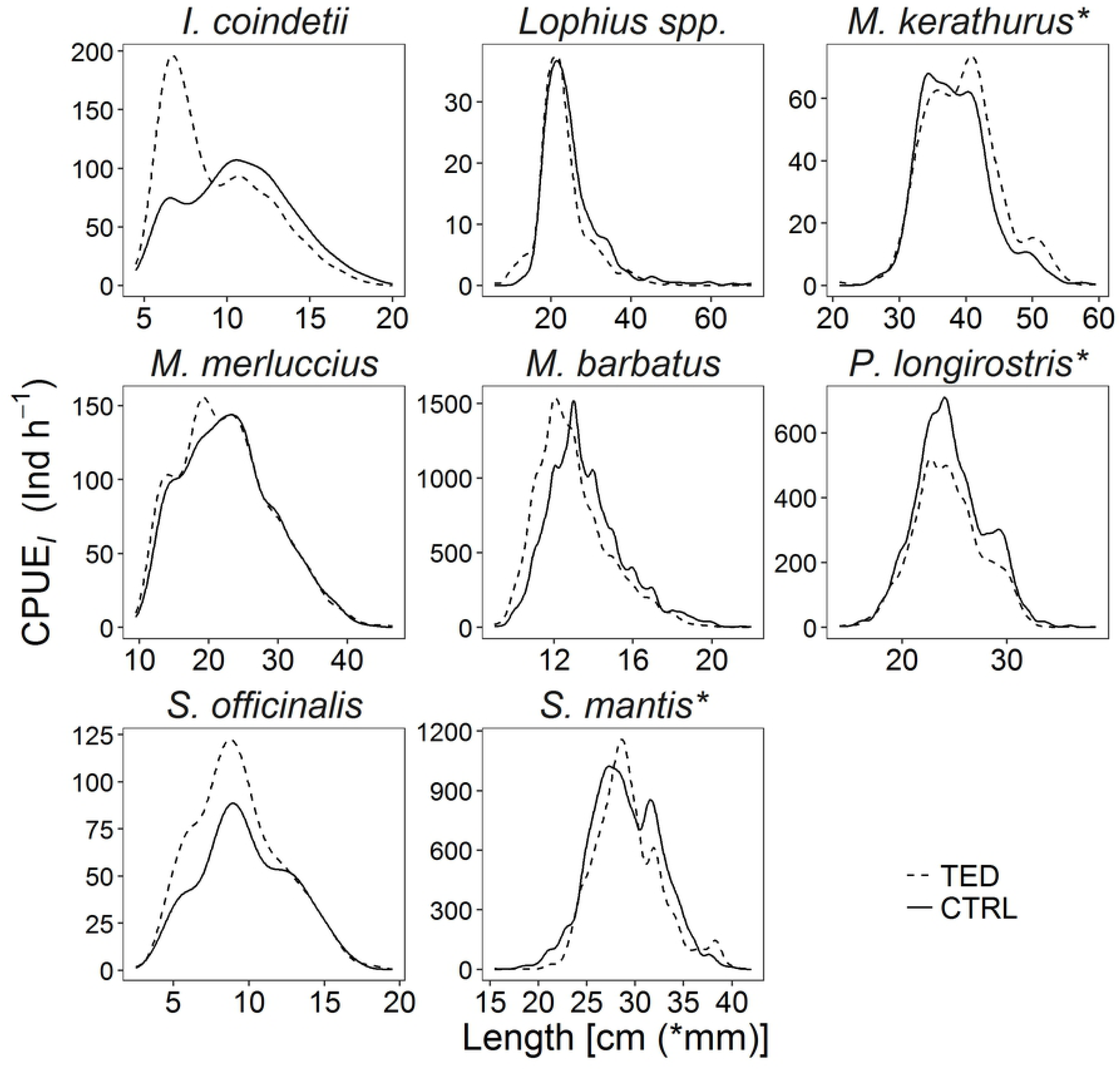
Pooled catch length-frequency distributions for each commercially important species for all boats.

**Fig. 6.**
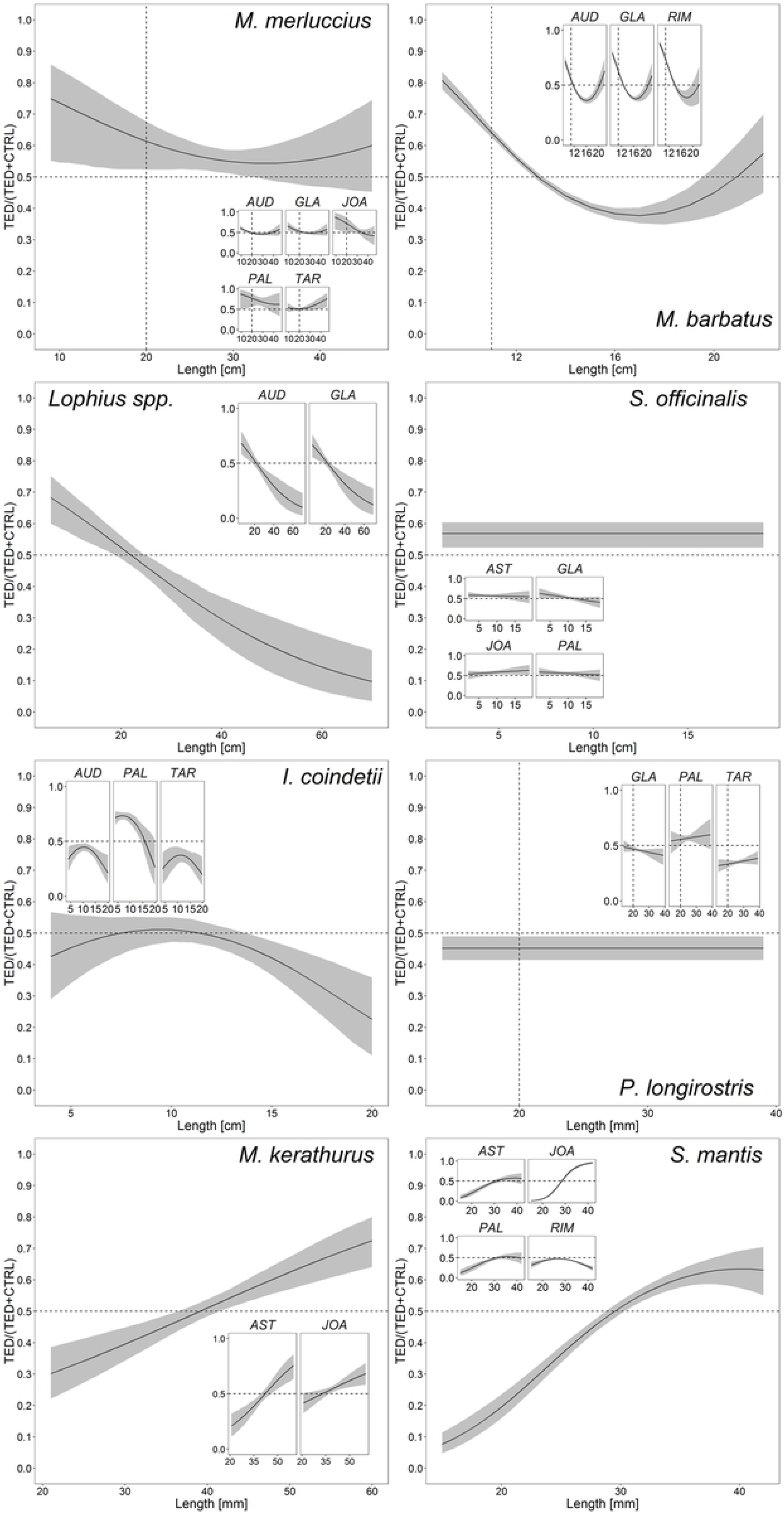
GLMM modelled proportions of the total catches caught by the TED. The main graphs represent the model for all boats together, while the smallest graphs represent the model applied to each single boat. Interpretation: a value of 0.5 indicates an even split between TED and CTRL, whereas a value of 0.25 indicates that the net mounting TED caught 25% of the total fish at that length and 75% were caught in the CTRL net. The shaded area is the 95% confidence interval.

Concerning the bycatch species, only *Pteroplatytrygon violacea* (3 individuals by JOA and 1 by PAL) and *Dasyatis pastinaca* (38 individuals caught by AST) represented bycatch of interest for conservation. Since the data were not enough no statistical analysis were performed. No turtles were caught during the trials, but several boats fishing in the same areas, reported the accidental catch of some individuals.

## Discussion

To our knowledge, this is the first experiment in the Mediterranean Sea to compare the catch efficiency between a traditional trawl and a trawl equipped with flexible TED, under professional operating conditions.

As already stated, in the Adriatic Sea the bottom trawlers mainly impact on the bulk of the sea turtle population, namely juvenile and sub-adult individuals. Consequently, the sea turtle population conservation mostly depends on the survival of the animal bycaught. Thus, the development of effective Bycatch Reducer Devices (BRDs) could be considered as an emergency to reduce the number of turtles bycaught.

In the present paper, the effectiveness of the TED compared to the CTRL net was demonstrated, as the weight and composition of the commercial part of the catch was not affected by the presence of the grid during fishing operations. On the other hand, the marine debris and litter were significantly reduced in the TED net, with the consequence of improving the quality of the catch by removal of large objects potentially damaging the catch itself. These results are in accordance with those obtained by [36]. Recently, [37] showed the persistence of several litter categories in the northern Adriatic. The authors observed the highest litter concentrations within 30 m depth and the lowest values between 30 and 50 m. Our findings on litter followed the same bathymetry distribution, with the highest concentrations in the shallower and medium hauls and the lowest deeper.

The LFDs of the main commercial species were similar between TED and CTRL; the performances of the two gears appeared similar without any significant loss of sizes, with the only exception for largest individuals of *Lophius* spp.. Moreover, Elasmobranches species such as rays and sharks were mostly entrapped in the CTRL nets. The results obtained with *Lophius* spp, sharks and the less presence of large marine litter could be considered as a proof of effectiveness of the TED also towards large animals, such as turtles. A slight depletion due to the presence of TED was observed for *M. barbatus, M. merluccius* and *I. coindetii*, even if without any significant difference. For crustaceans, on the contrary, a major effectiveness in catching of the TED. This could be due to the reduction in garbage for the TED. In fact, the litter can crush the fragile crustaceans in the traditional nets losing animals potentially marketable. From these results, TED appeared as a promising device to be implemented in the traditional gears without compromising the commercial catches as already stated by [36].

Opposite to the protection of sea turtle nesting sites, the measures taken to identify mitigation devices and strategies appropriate to mitigate the threat posed by fisheries are very low (see [38] for a review of the conservation measures). Conservation measures for sea turtles interacting with fisheries were reported in several papers and for several fishing gears [38]. The measures to reduce damages provoked by pelagic longlines to sea turtle include the use of “circle hooks”, difficult to be ingested, but, so far, the results are still controversial [39]. Concerning set nets, the only effective countermeasure to avoid the sea turtle tangle is the use of special lamps mounted on the net permitting the turtle to avoid them [40,41]. For the bottom trawls the only mitigation measure tested, so far, is the TED. The new flexible TED experimented by [36], was sufficiently stiffer and less flexible, than previous tested [22,42], to maintain the rigid configuration to the net, but flexible enough to winding safely around a standard net winch. This also translates into no changes of the on board procedures, or instruments, and with no loss of time during hauling.

The technical changes reported above are not mandatory in any of the Mediterranean countries, but they have been tested and promoted only on a voluntary basis or under economic incentives. Besides the technical innovations, other countermeasures to reduce fishery bycatch consist in informing on the results of the experimental researches, sensitize and training fishers on the best practices of sea turtle recovery after the catch. Indeed, one of the main problem to face is the reluctance of fishers in modifying the gears, something perceived as a reduction in profit, or increase fuel consumption. In this regard, the compliance with fishers is fundamental for bycatch reduction and depends on the incentives given to them [43]. If scientists will be able to demonstrate that a modification of the gear does not involve any modification of the commercial catch, probably most of the fishers could be interested in the change. There are examples worldwide, that this strategy works [44]. So far, in the Adriatic Sea the responses of the fishers involved in the experimentation are promising, giving the sensation that also in the Mediterranean Sea, fishers could collaborate towards the safeguard of sea turtles. Finally, the combination of education, outreach programs, and cooperative fisheries management, provide a model of participatory bycatch assessments and ultimately bycatch mitigation [45].

In conclusion, conservation of sea turtles over a wide area, as it is the Mediterranean Sea, is politically challenging. Moreover, the measures involving technical modifications of fishing gears require significant investments but are technically feasible and could guarantee the success of the conservation.

## Acknowledgements

The authors are indebted with the crews of the vessels for their support and with all the students and personnel of the CNR – IRBIM of Ancona that kindly helped with laboratory activities.

## Supporting information

**S1 Table. List of the commercial species caught during the trials with associated average CPUEW and standard errors.**

**S2 Table. List of the discard species caught during the trials with associated average CPUE**_**W**_ **and standard errors.**

**S3 Table. : List of the discard species caught during the trials with associated average CPUE**_**W**_ **and standard errors.** Marine litter codes as follow: A = PLASTIC: 01 = bottle; 02 = sheet; 03 = bag; 05 = fishing line (monofilament); 07 = synthetic rope; 08 = fishing net; 09 = cable ties; 10 = strapping band; 11 = crates and containers; 12 = mussel farming ropes; 13 = other. C = METAL: 01 = cans (food); 02 = cans (beverage); 03 = fishing related; 07 = cables; 08 = other. D = RUBBER; 01 = boots; 02 = balloons; 04 = tyre; 05 = glove; 06 = other. E = GLASS/CERAMIC: 01 = jar; 02 = bottle; 04 = other. F = NATURAL PRODUCT: 01 = wood (processed); 02 = rope; 05 = other. G = MISCELLANEUS: 01 = clothing/rags; 03 = other. Debris shell = empty shells; debris echinoderms = piece of sea urchins or dead sea urchins; debris wood = natural wood (branches or tree trunk); debris organic = unidentified organic material.

**S1 Fig. Fishes swimming in front of the TED and passing easily through its bars during trawling.**

**S2 Fig. Detail of an Angler fish (*Lophius* spp.) (inside the red round) blocked by the bars of the TED.** This could happen because of the particular shape of this species, characterized by a big head. In some instances, the individuals were pushed to enter the grid by the hydrodynamic force itself, while in other cases, the animals rolled up until to reach the opening before the TED on the upper side of the net. Although the TED is not the same of that used during the trials, the effect of the flexible TED’s bars on this species is the same.

## References

1. Bianchi CN, Morri C. Marine biodiversity of the Mediterranean Sea: Situation, problems and prospects for future research. Mar Pollut Bull. 2000;40: 367–376. doi:10.1016/S0025-326X(00)00027-8

2. Colloca F, Cardinale M, Maynou F, Giannoulaki M, Scarcella G, Jenko K, et al. Rebuilding Mediterranean fisheries: A new paradigm for ecological sustainability. Fish Fish. 2013;14: 89–109. doi:10.1111/j.1467-2979.2011.00453.x

3. Tudela S. Ecosystem effects of fishing in the mediterranean. An analysis of the major threats of fishing gear and practices to biodiversity and marine habitats. Stud Rev Gen Fish Comm Mediterr. Rome: FAO; 2004;74: 44. doi:10.1017/CBO9781107415324.004

4. Worm B, Barbier EB, Beaumont N, Duffy JE, Folke C, Halpern BS, et al. Impacts of biodiversity loss on ocean ecosystem services. Science (80-). 2006;314: 787–790. doi:10.1126/science.1132294

5. Bearzi G. Interactions between cetaceans and fisheries in the Mediterranean Sea. In: Notarbartolo di Sciara G, editor. Cetaceans of the Mediterranean and Black Seas: State of the knowledge and conservation strategies. Monaco: ACCOBAMS; 2002. p. 92.

6. Casale P. Sea turtle by-catch in the Mediterranean. Fish Fish. 2011;12: 299–316. doi:10.1111/j.1467-2979.2010.00394.x

7. Ferretti F, Myers RA, Serena F, Lotze HK. Loss of large predatory sharks from the Mediterranean Sea. Conserv Biol. 2008;22: 952–964. doi:10.1111/j.1523-1739.2008.00938.x

8. European Commission. Council Directive 92/43/EEC of 21 May 1992 on the conservation of natural habitats and of wild fauna and flora. Council of the European Communities (CEC). Off J Eur Communities. 1992;206: 7–50.

9. FAO. International Guidelines on bycatch management and reduction of discards. [Internet]. FAO International Guidelines. 2011. doi:ISSN 2070-6987

10. Lewison RL, Johnson AF, Verutes GM. Embracing Complexity and Complexity-Awareness in Marine Megafauna Conservation and Research. Front Mar Sci. 2018;5: 1–11. doi:10.3389/fmars.2018.00207

11. Casale P, Tucker AD. Caretta caretta. IUCN Red List Threat Species 2015. 2015;8235: 20. doi:http://dx.doi.org/10.2305/IUCN.UK.2015-4.RLTS.T3897A83157651.en

12. Casale P. Caretta caretta (Mediterranean subpopulation). IUCN Red List Threat Species 2015. 2015; e.T83644804A83646294.

13. Lucchetti A, Vasapollo C, Virgili M. An interview-based approach to assess sea turtle bycatch in Italian waters. PeerJ. 2017;5: e3151. doi:doi.org/10.7717/peerj.3151

14. Lucchetti A, Sala A. An overview of loggerhead sea turtle (Caretta caretta) bycatch and technical mitigation measures in the Mediterranean Sea. Rev Fish Biol Fish. 2010;20: 141–161. doi:10.1007/s11160-009-9126-1

15. Casale P. Incidental catch of marine turtles in the Mediterranean Sea: Captures, mortality, priorities. Rome, Italy: WWF; 2008.

16. Lucchetti A, Pulcinella J, Angelini V, Pari S, Russo T, Cataudella S. An interaction index to predict turtle bycatch in a Mediterranean bottom trawl fishery. Ecol Indic. Elsevier Ltd; 2016;60: 557–564. doi:10.1016/j.ecolind.2015.07.007

17. Lazar B, Margaritoulis D, Tvrtkovic N. Tag recoveries of the loggerhead sea turtle Caretta caretta in the eastern Adriatic Sea: implications for conservation. J Mar Biol Assoc UK. 2004;84: 475–480. doi:10.1017/S0025315404009488h

18. Casale P, Laurent L, De Metrio G. Incidental capture of marine turtles by the Italian trawl fishery in the north Adriatic Sea. Biol Conserv. 2004;119: 287–295. doi:10.1016/j.biocon.2003.11.013

19. Epperly S. Fisheries-related mortality and turtle excluder devices (TEDs). In: Lutz P, Musick J, editors. The biology of sea turtles. Boca Raton, FL: CRC Press; 2003. pp. 339–353.

20. Lewison R, Wallace B, Alfaro-Shigueto J, Mangel JC, Maxwell SM, Hazen EL. Fisheries bycatch of marine turtles: lessons learned from decades of research and conservation. In: Wyneken J, Lohmann KJ, Musick JA, editors. Biology of sea turtles, vol 3. Boca Raton, FL: CRC Press; 2013. pp. 329–351.

21. Atabey S, Taskavak E. A preliminary study on the prawn trawls excluding sea turtles. J Fish Aquat Sci. 2001;18: 71–79.

22. Sala A, Lucchetti A, Affronte M. Effects of Turtle Excluder Devices on bycatch and discard reduction in the demersal fisheries of Mediterranean Sea. Aquat Living Resour. 2011;24: 183–192. doi:10.1051/alr/2011109

23. Mitchell J, Watson J, Foster D, Caylor R. The Turtle Excluder Device (TED): a guide to better performance. 1995.

24. Eayrs S. A Guide to Bycatch Reduction in Tropical Shrimp-Trawl Fisheries. Fao. 2007; 124. Available: http://www.fao.org/docrep/field/003/ab825f/AB825F00.htm#TOC

25. Quinn G, Keough M. Experimental Design and Data Analysis for Biologists. Cambridge University Press; 2002.

26. Zuur A, Ieno E, Walker N, Saveliev A, Smith G. Mixed effects models and extensions in ecology with R. New York: Springer; 2009.

27. Pinheiro J, Bates D. Mixed-Effects Models in S and S-PLUS. Springer, editor. 2000.

28. Holst R, Revill A. A simple statistical method for catch comparison studies. Fish Res. 2009;95: 254–259. doi:10.1016/j.fishres.2008.09.027

29. Fryer RJ, Zuur AF, Graham N. Using mixed models to combine smooth size-selection and catch-comparison curves over hauls. Can J Fish Aquat Sci. 2003;60: 448–459. doi:10.1139/f03-029

30. Fryer RJ. A model of between-haul variation in selectivity. ICES J Mar Sci. 1991;48: 281–290. doi:10.1093/icesjms/48.3.281

31. Van Marlen B, Wiegerinck JAM, van Os-Koomen E, van Barneveld E. Catch comparison of flatfish pulse trawls and a tickler chain beam trawl. Fish Res. Elsevier B.V.; 2014;151: 57–69. doi:10.1016/j.fishres.2013.11.007

32. Browne D, Minto C, Cosgrove R, Burke B, McDonald D, Officer R, et al. A general catch comparison method for multi-gear trials: Application to a quad-rig trawling fishery for Nephrops. ICES J Mar Sci. 2017;74: 1458–1468. doi:10.1093/icesjms/fsw236

33. R Core Team. R: A language and environment for statistical computing. [Internet]. R Foundation for Statistical Computing. Vienna; 2016. doi:3-900051-14-3

34. Pinheiro J, Bates D, DebRoy S, Sarkar D. nlme: Linear and Nonlinear Mixed Effects Models. 2018.

35. Bates D, Martin M, Bolker B, Walker S. Fitting Linear Mixed-Effects Models Using lme4. J Stat Softw. 2015;67: 1–48.

36. Lucchetti A, Punzo E, Virgili M. Flexible Turtle Excluder Device (TED): an effective tool for Mediterranean coastal multispecies bottom trawl fisheries. Aquat Living Resour. 2016;29: 1–12. Available: http://10.0.4.27/alr/2016016

37. Strafella P, Fabi G, Despalatovic M, Cvitković I, Fortibuoni T, Gomiero A. Assessment of seabed litter in the Northern and Central Adriatic Sea (Mediterranean) over six years. Mar Pollut Bull. Elsevier; 2019;141: 24–35. doi:10.1016/j.marpolbul.2018.12.054

38. Casale P, Broderick AC, Caminas JA, Cardona L, Carreras C, Demetropoulos A, et al. Mediterranean sea turtles: current knowledge and priorities for conservation and research. Endanger Species Res. 2018;36: 229–267. doi:10.1136/bmj.b2700

39. Piovano S, Basciano G, Swimmer Y, Giacoma C. Evaluation of a bycatch reduction technology by fishermen: a case study from Sicily. Mar Policy. Elsevier; 2012;36: 272–277. doi:10.1016/j.marpol.2011.06.004

40. Ortiz N, Mangel JC, Wang J, Alfaro-Shigueto J, Pingo S, Jimenez A, et al. Reducing green turtle bycatch in small-scale fisheries using illuminated gillnets: the cost of saving a sea turtle. Mar Ecol Prog Ser. 2016;545: 251–259. doi:10.3354/meps11610

41. Virgili M, Vasapollo C, Lucchetti A. Can ultraviolet illumination reduce sea turtle bycatch in Mediterranean set net fisheries? Fish Res. 2018;199: 1–7. doi:10.1016/j.fishres.2017.11.012

42. Lazar B, Tvrtkovic N. Marine turtles in the eastern part of the Adriatic sea: preliminary research. Nat Croat. 1995;4: 59–74.

43. Cox TM, Lewison RL, Žydelis R, Crowder LB, Safina C, Read AJ. Comparing effectiveness of experimental and implemented bycatch reduction measures: The ideal and the real. Conserv Biol. 2007;21: 1155–1164. doi:10.1111/j.1523-1739.2007.00772.x

44. Finkbeiner EM, Wallace BP, Moore JE, Lewison RL, Crowder LB, Read AJ. Cumulative estimates of sea turtle bycatch and mortality in USA fisheries between 1990 and 2007. Biol Conserv. Elsevier Ltd; 2011;144: 2719–2727. doi:10.1016/j.biocon.2011.07.033

45. Hall M, Nakano H, Clarke S, Thomas S. Working with fishers to reduce bycatches. In: Kennelly S, editor. Bycatch Reduction in the World’s Fisheries. Springer-Verlag; 2007.

